# Phase-Amplitude Coupling Detection and Analysis of Human 2-Dimensional Neural Cultures in Multi-well Microelectrode Array in Vitro

**DOI:** 10.1101/2023.07.08.548227

**Authors:** Yousef Salimpour, William S. Anderson, Raha Dastyeb, Shiyu Liu, Guo-li Ming, Hongjun Song, Nicholas J. Maragakis, Christa W. Habela

**Affiliations:** Functional Neurosurgery Laboratory, Department of Neurosurgery, Johns Hopkins School of Medicine, Baltimore, Maryland, USA; Department of Neurology, Johns Hopkins School of Medicine, Baltimore MD, USA; Department of Neuroscience and Mahoney Institute for Neurosciences, Perelman School of Medicine, Philadelphia, PA, USA

**Author notes:** Correspondence: Yousef Salimpour, MSc, PhD, Christa W. Habela MD. PhD. Co-Senior Authors (Nicholas J. Maragakis, Christa W. Habela). **Author contributions:** YS: performed analysis, wrote first draft, conceptualized experiments and contributed to the final draft of the manuscript, WA: conceptualized experiments and contributed to the final draft of the manuscript. RD contributed to analysis and writing. SL contributed to experiments. HS: contributed materials and to experimental protocols and analysis and writing. G-LM: contributed materials and to experimental protocols and analysis and writing. NM: conceptualized experiments and contributed to the final draft of the manuscript, provided materials and supervised the project. CWH performed experiments, analysis and wrote first draft of manuscript, conceptualized experiments and contributed to the final draft of the manuscript, provided materials and supervised the project.

**Keywords:** Cross-frequency coupling, phase-amplitude coupling, Multielectrode Array

## Abstract

Human induced pluripotent stem cell (hiPSC)_-_ derived neurons offer the possibility of studying human-specific neuronal behaviors in physiologic and pathologic states *in vitro*. However, it is unclear whether these cultured neurons can achieve the fundamental network behaviors that are required to process information in the human brain. Investigating neuronal oscillations and their interactions, as occurs in cross-frequency coupling (CFC), is potentially a relevant approach. Microelectrode array culture plates provide a controlled framework to study populations of hiPSC-derived cortical neurons (hiPSC-CNs) and their electrical activity. Here, we examined whether networks of two-dimensional cultured hiPSC-CNs recapitulate the CFC that is present in networks *in vivo*. We analyzed the electrical activity recorded from hiPSC-CNs grown in culture with hiPSC-derived astrocytes. We employed the modulation index method for detecting phase-amplitude coupling (PAC) and used an offline spike sorting method to analyze the contribution of a single neuron’s spiking activities to network behavior. Our analysis demonstrates that the degree of PAC is specific to network structure and is modulated by external stimulation, such as bicuculine administration. Additionally, the shift in PAC is not driven by a single neuron’s properties but by network-level interactions. CFC analysis in the form of PAC explores communication and integration between groups of nearby neurons and dynamical changes across the entire network. *In vitro*, it has the potential to capture the effects of chemical agents and electrical or ultrasound stimulation on these interactions and may provide valuable information for the modulation of neural networks to treat nervous system disorders *in vivo*.

**Significance:** Phase amplitude coupling (PAC) analysis demonstrates that the complex interactions that occur between neurons and network oscillations in the human brain, *in vivo*, are present in 2-dimensional human cultures. This coupling is implicated in normal cognitive function as well as disease states. Its presence *in vitro* suggests that PAC is a fundamental property of neural networks. These findings offer the possibility of a model to understand the mechanisms and of PAC more completely and ultimately allow us to understand how it can be modulated *in vivo* to treat neurologic disease.

## 1 Introduction

Our understanding of human neurologic development and disease is confounded by the fact that the study of human neurons *in vivo* is limited by patient safety factors. Invasive intracranial recordings used in the diagnosis and management of neurologic disorders, such as epilepsy, allow significant observations to be made about human neural network behaviors.

However, there are limited opportunities to test hypotheses about the mechanisms and clinical implications of this neural activity, specifically when it comes to single-cell activity that may underlie network changes. Computational models have been the mainstay of hypothesis testing, but without understanding the underlying behavior and biology of single cells that generate this activity, these models are imperfect ^1^.

In human neurologic disorders, such as epilepsy and Parkinson’s disease (PD), cross-frequency coupling (CFC), which describes the interactions of neural oscillations at different frequencies with one another, is one parameter to describe network behavior. The presence or absence of CFC has been implicated as an important feature of functional neural network activity based on an association with both normal processing behaviors and pathologic states, such as seizures ^2^. CFC is present and increased at the onset of seizures, but not during or in between seizures ^3^. It is also present and increased during complex cognitive processing ^4^. Phase-amplitude coupling (PAC), a form of cross frequency coupling, describes the relationship between the phase of low frequency oscillations and the amplitude of high frequency activity. In the motor cortex of PD patients, excessive abnormal PAC between the phase of beta rhythms and the amplitude of high frequency gamma activity has been demonstrated and strong correlations between PAC and motor symptom severity have been observed ^5^. Further, modulation of motor symptoms with levodopa or deep brain stimulation has been associated with a reduction in this PAC ^6^. This suggests that not only is PAC a biomarker for disease states, but that modulation of PAC may modulate symptoms. However, the underlying cellular processes contributing to this coupling in normal and pathologic states are not fully understood.

Human induced pluripotent stem cell (hiPSC)-derived neuronal cultures offer one potential platform to model and manipulate the network interactions of human neurons *in vitro*. In the 15 years since the original description of neural cell generation from patient-derived hiPSCs, numerous subsets of neurons and glial cells have been described and applied to the study of neurologic diseases ranging from neural developmental disorders to neurodegenerative disorders ^7 8^. The advantages of these systems include an intact human genetic background with the ability to examine spontaneously as well as induced genetic variation, high-throughput platforms allowing for the study and manipulation of these processes, and the ability to manipulate cell types and cell numbers within a network ^9^.

The use of hiPSCs in both two-dimensional monolayer cultures and three-dimensional “organoid” cultures has revealed that cultured neurons generate spontaneous spikes and bursting activity, indicating functional and excitable neurons as well as synchronous network activity indicative of synoptically interconnected neurons in functional networks ^10 9^. More complex network behaviors such as cross-frequency coupling have not been previously described in 2D (2 Dimensional) cultures, but the presence of this activity *in vitro* in 2D cultures would offer an intermediate step between computational modeling and *in vivo* applications to generate and test hypotheses. We hypothesize that PAC is a fundamental and intrinsic property of human neuronal networks and therefore predict that it can be detected in human neurons *in vitro* independent of three-dimensional structure or external inputs.

## 2 Materials and Methods

### Cell Culture

hiPSCs were maintained and passaged as previously described ^11^. Neural progenitor cells (NPCs) were generated using the human forebrain differentiation technique^12^. For differentiation to human cortical neurons, NPCs were thawed onto poly-L-ornithine (PLO) and laminin-coated plates into Neuronal media consisting of Neurobasal Media (Thermofisher), Glutamax (Thermofisher), nonessential amino acids (Thermofisher), B27 supplement (Thermofisher), BDNF (10ng/mL), GDNF (10ng/mL) supplemented with 125nM of notch inhibitor Compound E (1:2000 DMSO) and 20 μM of Rho-associated coiled-coil forming protein serine/threonine kinase inhibitor (ROCK-I, compound Y-27632) (1:500 DMSO). After 48 hours, ROCK-I was removed, and Neuronal media supplemented with Compound E (125nM) was completely exchanged every 48 to 72 hours until DIC 37 with every other media change containing 1 ug/mL laminin (Thermofisher). For astrocyte differentiation, NPCs were thawed onto PLO and laminin-coated plates and cultured in astrocyte media supplemented with 1% knockout serum replacement (KSR)(Thermofisher). Astrocyte progenitor cells (APCs) were passaged every 4 to 7 days with media changes every 48 to 72 hours. With the first passage, APCs were plated onto 1% Matrigel (Corning) coated plates and maintained on Matrigel throughout subsequent passages. In preparation for co-cultures, APCs aged to DIC 77 - 86 were thawed onto PLO and laminin-coated plates in astrocyte media 1% KSR until DIC 90.

### Combined astrocyte and neuron co-culture

Neurons were aged to DIC 37 and synchronized with DIC 90-95 APCs. Cells were washed with sterile phosphate-buffered saline and lifted using 0.05% Trypsin (Thermofisher) incubation at 37°C for 3 minutes. Neurons were resuspended for a final concentration of 50,000 cells per 5 μL in Neuronal media supplemented with 1% anti-anti (GIBCO), 5% KSR, 1:2000 compound E (MEA (Multielectrode Array) media) and 1:500 ROCK-I. Astrocytes were diluted to 25,000 cells per 5 μL in the same media as neurons. Neurons and astrocytes were combined 1:1 by volume. For Multi-electrode array (MEA) recordings, 5 μL of the neuron and astrocyte suspension was placed as a droplet onto the center of a PLO and laminin coated electrode array well and incubated 37°C, 5% CO_2_ for 20-30 minutes at before 500 μL of MEA media + ROCK-I was pipetted down the side of the well in two parts. ROCK-I was removed from the media after 48 hours and cocultures underwent partial media exchanges with MEA KSR twice a week until neuronal DIC 80 and subsequently once a week, with 1ug/ml laminin added every other media change.

### MEA Recordings

Plates were equilibrated in the Axion Edge at 37°C, 5% CO_2_ for a minimum of 3 minutes prior to recording. Five-minute recordings were obtained at weekly or bi-weekly intervals beginning on DIC 42 until DIC 218. For drug applications, recordings consisted of 3-minute baseline recordings followed by vehicle and then drug application while recording continued. Raw data was exported to MATLAB format for CFC detection analysis.

### Spike detection and sorting

To investigate the association between PAC and single cell spiking, we used an offline and unsupervised algorithm for spike detection and sorting using wavelets and super-paramagnetic clustering ^13^. Recording from each individual electrode fed to two parallel pipelines, one for spike detection and sorting and the other for PAC analysis.

### Instantaneous Phase, Frequency, and Amplitude Analysis

Raw voltage data from MEA recordings were analyzed using bandpass filtering in conjunction with the Discrete Hilbert Transform (DHT) to extract phase data from low frequency and amplitude from high frequency activity ^14^. The raw signals were filtered in the high-frequency (20-200 Hz) and low-frequency (1-10 Hz) frequency bands using a zero-phase bandpass filter. Subsequently, the DHT was applied, which yielded an analytical function. The phase and absolute value of the analytic function from the filtered recording was used to calculate the instantaneous phase, frequency and amplitude.

### Low-frequency Phase Modulated High-frequency Power

Phase amplitude coupling measures the degree to which the high-frequency power is modulated by the phase of the low-frequency oscillations. To quantify PAC, the mean modulation index (MMI) was used. MMI is defined based on modulation index (MI) estimation ^14-16^ which is summarized as follows: For each time window, the average amplitude of the high-frequency signal within that window is calculated. Next, the phase (the position or angle of the low-frequency signal within its cycle) of the low-frequency signal at the center of each time window is determined. Then the amplitude of the high-frequency signal within each time window is compared to the overall average amplitude of the signal to determine if the amplitude is larger or smaller than the average. Finally, the modulation index is calculated by averaging the difference between the amplitude values and the average amplitude. A higher modulation index indicates a stronger coupling between the phase and amplitude signals, suggesting that the phase of the low-frequency signal has a greater impact on the amplitude of the high-frequency signal ^6,17^.

### Statistical Analysis

The MATLAB Statistical package was used. To assess the statistical significance of PAC changes, each experimental testing section, including a period during the retrieval phase, was analyzed using surrogate control analysis (Tort et al., 2010). We performed a statistical control analysis on each session by creating shuffled versions of the complex time series consisting of low-frequency phase and high-frequency amplitude. This generated surrogate MI values, from which the MI chance distribution could be inferred ^14^. One-sample and paired-sample t-test tests were used for mean MI changes, and a *P* < 0.05 was considered statistically significant

## 3 Results

### Phase amplitude coupling *in vitro*

Human iPSC-derived human cortical neurons (hiPSC-CNs) in co-culture with hiPSC-derived human cortical astrocytes were grown in multi-well MEA plates. Continuous voltage data from recordings on DIC 218 showed that CFC exists in the form of PAC in two-dimensional cultures (Figure 1). The raw voltage recording from a single representative contact (Figure 1A, gray trace) was subject to zero phase band pass filters in the low-frequency range of 1-10 Hz (Figure 1A, red trace and high-frequency range of 20-200 Hz (Figure 1B, teal trace). Visual inspection of these traces suggests that the peak amplitude of the high frequency component is related to the phase of the low frequency oscillation (Figure 1B). To determine whether there is in fact a coupling between low frequency phase and the amplitude of high frequency activity, the Hilbert transformation ^14^ was used. The phase values of low frequency activity (Figure 1C, red trace) and the power of high frequency components (Figure 1C, teal trace) were computed from the corresponding raw voltage over time in Figure 1B.. The time – frequency representation was used to set the range for exploring PAC ^18^.. Figure 1D demonstrates the frequency component of the time segment from figure 1B and demonstrates a wide range of frequency components in the raw data.

**Figure 1.**
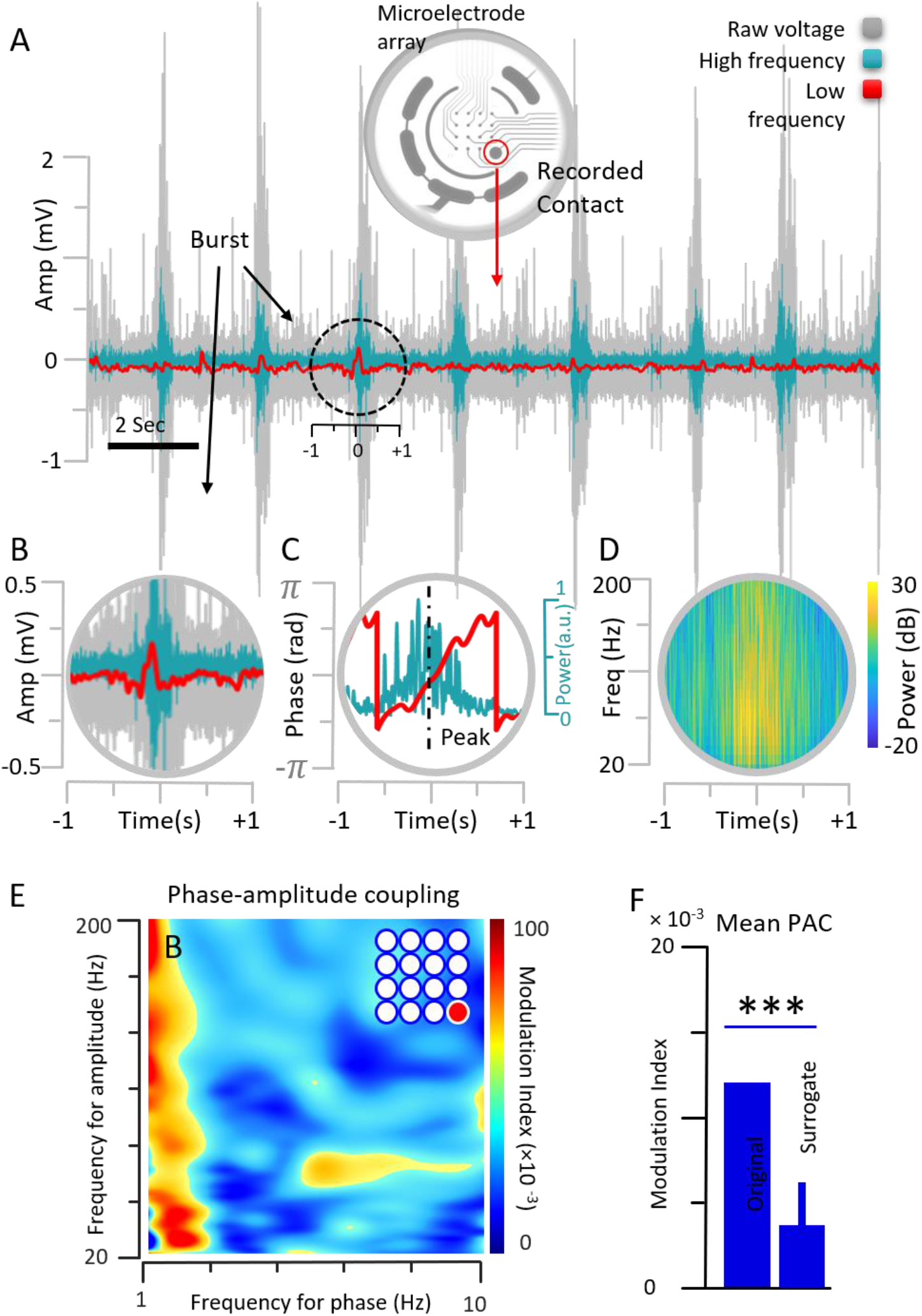
Frequency analysis in Multi-electrode Array Recordings. **A**. Sample recording from a single electrode of a microelectrode array is shown. Raw local field potential is grey. Zero phase bandpass filtering of the recorded data in low frequency range (1-10Hz) in red and high frequency range (20-200Hz) in teal. **B**. Amplification of the dashed circle in A. Visual inspection of these traces suggests a typical signature of cross-frequency coupling (CFC) in the format of the phase-amplitude coupling (PAC). **C**. Phase decomposition of the low frequency (red) and power decomposition of the high frequency range (teal) corresponding to the voltage traces in B shows the association between the peak of the phase in low frequency (dashed line) and the power of high frequency components. **D**. The time – frequency representation shows a wide range of frequency components in the raw voltage data associated with sample burst demonstrated in panel B. **E**. The modulation index measure is used to quantify and visualize PAC in the same electrode. **C**. Significant differences between estimated mean PAC and a set of randomly created shuffled versions of the low-frequency phase and high-frequency amplitude (0.0039 ± 0.0022, *P* < 0.0001).

To quantify PAC, the modulation index-based measurement was used ^14^. The results reveal a prominent level of PAC between the phase of low frequency activity (1 – 10 Hz) and the amplitude of high frequency activity (20-200 Hz) (Figure 1E). In this example, the coupling is most pronounced in the low frequency range 1-2 Hz (highest intensity on heat map) (Figure 1E). To verify the validity of PAC detection, a surrogate control analysis was used. This was accomplished by generating a new series of data that comes from maintaining the phase of the low frequency and shuffling the amplitude of the high frequency and estimating the modulating index based on these regenerated time series. This was replicated 10 times resulting in a significantly lower mean modulation index in the surrogate than is present in the original (one-sample t-test *P* < 0.0001) (Figure 1F). This excludes a random origin of the observed PAC detection in our cultures ^14^.

### Cross-frequency across the network

To further validate the observed phenomenon of PAC in monolayer culture *in vitro*, additional electrodes from the same culture well shown in Figure 1 were analyzed. Three additional representative electrodes are shown (Figure 2A, top and bottom left). When surrogate analysis is compared to the PAC across 16 electrodes, there is a significant difference between surrogate (0.0064 ± 0.0024) and original measures (0.0136 ± 0.0049) (*P* < 0.0001) (Figure 2B). This demonstrates that CFC in the form of PAC is not limited to a single electrode and is detectable on all electrodes in the well. Additionally, similarities in PAC at different contacts suggests that the phenomenon is a network phenomenon and not dependent on the cells located at that electrode. When additional wells were analyzed (Figure 2C and D), the data indicate that this phenomenon is not unique to a particular well although there is some variation between wells.

**Figure 2.**
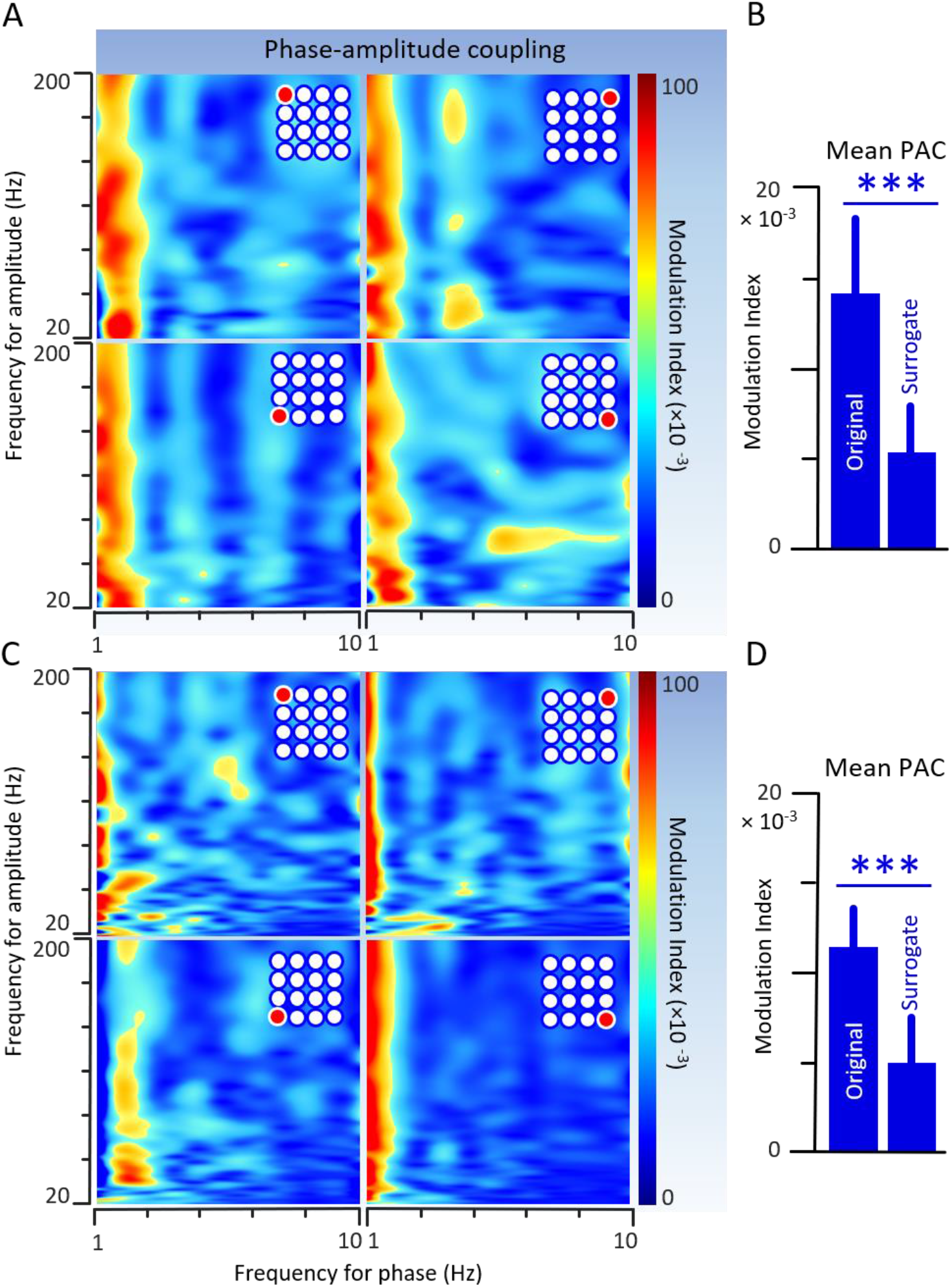
Cross frequency coupling (CFC) at the Network Level. CFC in the form of phase-amplitude coupling detected across an entire well. **A**. Three additional electrodes from the same culture well in **Figure 1** are shown. **B**. The mean PAC measures across the entire well are compared to the mean of the surrogate data. When surrogate analysis is compared to the PAC across 16 electrodes, there is a significant difference between surrogate (0.0064 ± 0.0024) and original measures (0.0136 ± 0.0049) (*P* < 0.0001). **C**. The results of PAC analysis from an additional well are shown here, indicating this phenomenon is not unique to a particular well. **D**. Similar to the representative well, the mean PAC measures across the entire well are compared to the mean of the surrogate data and there is still a significant difference between surrogate (0.0046±0.0029) and original measures (0.0111 ± 0.0025) (*P* < 0.0001).

### Detection of dynamic changes using CFC

CFC detection and analysis can be considered an effective tool to assess the dynamic changes of cultured neurons at a network level. To evaluate whether dynamic changes can be captured in vitro using PAC, we analyzed baseline activity as well as the changes occurring with the addition of the Gama-aminobutyric acid (GABA) receptor antagonist, bicuculine. Application of 10 μM bicuculine results in changes in the raw multiunit neuronal firing patterns (Figure 3A). When the period prior to drug application (left) is compared to the period after drug application (right), a change in the PAC is evident with a shift in the frequency of the low frequency driving the high frequency (Figure 3B). Dynamic changes in PAC over time can be demonstrated by sliding windows based dynamic PAC analysis ^6,17^ in which the high frequency amplitude across all low frequency amplitudes is analyzed over time. Applying this analysis to the raw data in Figure 3A, demonstrates that at the time of bicuculine administration there is a change in the PAC characterized by a transient increase in coupling followed by a decrease in coupling (Figure 3C), consistent with dynamic fluctuations in synchronization.

**Figure 3.**
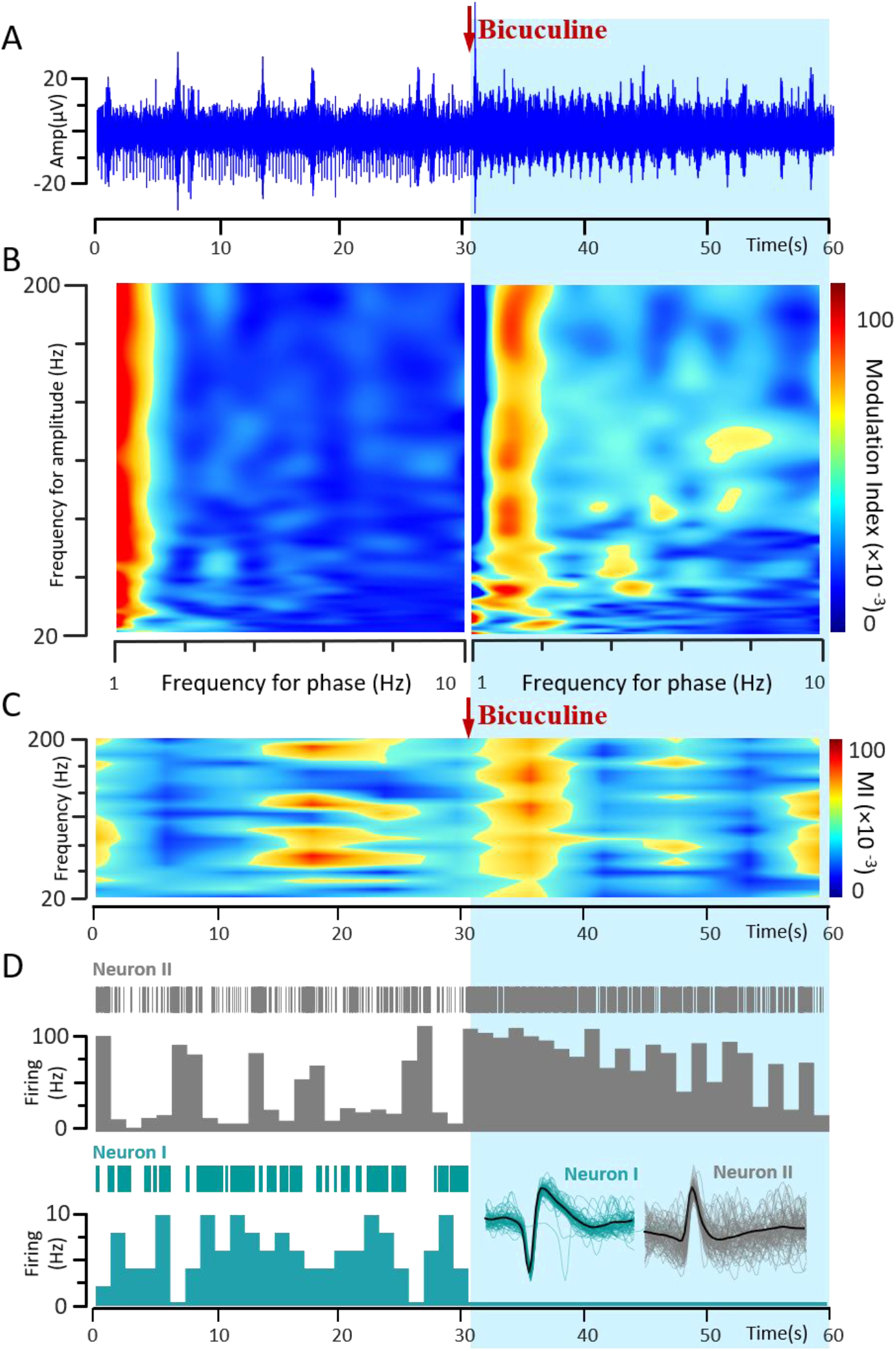
Cross frequency coupling (CFC) Reflection at the single cell level. **A**. Application of bicuculine (10 μM), indicated by red arrow and blue shading, results in changes in the raw multi-unit neuronal firing patterns. **B**. Modulation index-based quantification of phase-amplitude coupling show strong coupling between phase of the low frequency rhythm and high frequency oscillation which could be altered by bicuculine. **C**. Dynamic changes in phase amplitude coupling (PAC) over time can be demonstrated by sliding windows based dynamic PAC analysis in which the high frequency amplitude across all low frequency amplitudes is analyzed over time. **D**. Spike sorting of the local field potential recording from a single electrode reveals the association between CFC changes and single neuron firing pattern. Modulation index-based PAC measurement shows the reflection of CFC in excitatory and inhibitory neurons in the neuronal network. At the single-cell level, bicuculine application results in an increase in the firing rate and more tonic activity in some neurons (top) while other neurons have significantly reduced firing or stop firing (bottom). Raster plots (top) and spike rates (bottom) are displayed for example Neuron I (grey) and II (teal). Sorted spike wave forms for both neurons are displayed in the bottom right.

To study the relationship between dynamic network activity and single cell activity, offline spike sorting of the raw voltage data was performed ^13^. At the single-cell level, bicuculine application results in an increase in the firing rate and more tonic activity in some neurons (Figure 3D, top), while other neurons have significantly reduced firing or stop firing all together (Figure 3D, bottom). When these single cell changes are compared to the PAC changes, the changes in individual neuronal firing patterns outlast the transient increase in PAC, suggesting that the observed change in PAC is not reflective of single neuron properties but of the interactions between individual neurons at the network level.

## 4 Discussion

In this study, we investigated the formation of phase-amplitude coupling (PAC) in hiPSC-derived human cortical neurons in 2D cultures *in vitro*. The results demonstrate that, like the human brain *in vivo*, cross frequency coupling (CFC) is present in the form of PAC. This coupling is a property of the network of individual cultures with similar PAC across different electrodes in the same well and varying degrees of PAC in different wells. Additionally, the presence of PAC is dynamic and can be modulated by chemical manipulation of neuronal synaptic interactions. Finally, simultaneous analysis of single cell properties and PAC demonstrates that changes in PAC are not driven by the single cell properties of one neuron but rather the interactions between neurons.

Neuronal oscillations of different frequencies in the cortical and subcortical structures demonstrate complicated coupling properties reflecting more complex features of neural oscillations present in the brain’s normal activities, mostly in CFC in the form of PAC ^4^. CFC formation in hiPSC based neuronal cultures supports the theory that complicated neuronal oscillations are inherent properties of neuronal networks. The presence of PAC in human neuronal culture *in vitro* suggests that the intrinsic interactions between individual cells are sufficient to produce a specific level of local PAC without external influences. Phase amplitude coupling between delta oscillations has been described in 3-dimensional brain organoid cultures ^19^ and neuronal spiking in brain organoids has also been shown to be phase locked to theta oscillations ^20^. However, the phenomenon of PAC has not previously been described in 2D human cultures. The existence of PAC in 2D cultures suggests that PAC is a fundamental property of neural networks and is independent of 3D structure. Importantly, the presence of this phenomenon *in vitro* offers the possibility of a model system that can be used to further define the underlying determinants of PAC and its dynamics. Compared to 3D cultures, 2D cultures may provide specific advantages for future exploration including the ability to titrate the ratio of inhibitory to excitatory cells, genetically manipulate different cell types within the same cell culture system or examine the effects of non-neuronal cells such as astrocytes on PAC.

Although the neural mechanism underlying PAC in the brain is not fully understood, current evidence shows the contribution of both pyramidal cells and interneurons, which fire simultaneously at specific phases of low-frequency oscillation ^21^. The culture technique used here generates glutamatergic forebrain specific neurons with 5-10% GABAergic neurons ^12^. The change in multiunit and single unit firing rates in response to application of the GABA receptor antagonist, bicuculine demonstrates that there are functional GABAergic synapses in these networks. Importantly, the shift in the frequency for phase of the PAC in response to bicuculine confirms that synaptic GABAergic activity modulates the coupling. However, the fact that the change does not correlate with a simple increase or decrease in firing rate of individual neurons indicates that this phenomenon reflects network processing. *In vitro* 2D culture systems are potentially ideal model systems to study the contributions of both excitatory and inhibitory neurons to this process. Our evidence demonstrates the potential of CFC analysis to capture the dynamics of neuronal interactions at the network level and potentially explore the effects of different neuromodulation modalities such as chemical, electrical, and even ultrasound in the future.

HiPSC-derived neurons offer the potential for the development of cell therapies, drug discovery, and disease modeling. However, the extent to which *in vitro* populations of neurons can recapitulate the complex network interactions that are present in the brain *in vivo* is not fully understood. Disruption of these interactions, that is not necessarily evident in single-cell properties, may be the underlying pathology in numerous disease models such as epilepsy and PD. Application of neuronal network analysis in hiPSC-derived neuronal cultures provides a unique opportunity for investigating the neuronal mechanisms underlying complicated behavior of neuronal populations. It adds another dimension to the analysis of hiPSC-derived neurons and provides the potential to enhance our understanding of complex oscillatory behavior in neuronal information processing.

## Acknowledgements

CNCDPK12, K08NS102526, Doris Duke Foundation Clinical Scientist Award to CWH; NIH 5R01NS117604 and Department of Defense ALSRP W81XWH1810175 to N.J.M.; NIH R35NS116843 to H.S. and NIH R35NS097370 to G-l.M

